# Testing *Anopheles* larvae and adults using standard bioassays reveals susceptibility to chlorfenapyr (pyrrole) while highlighting variability between species

**DOI:** 10.1101/2024.03.24.586483

**Authors:** Calmes Bouaka, Marilene Ambadiang, Fred Ashu, Caroline Fouet, Colince Kamdem

## Abstract

A standard test is available for assessing the susceptibility of adult *Anopheles* mosquitoes to chlorfenapyr, a new active ingredient in insecticide-treated nets. However, for a new insecticide with a unique mode of action, testing both larvae and adults using different routes of exposure is crucial to a comprehensive evaluation of susceptibility and to identifying potential selection pressures that may drive resistance. We followed WHO guidelines to assess the lethal toxicity of chlorfenapyr and monitor *Anopheles* susceptibility. Based on the median lethal concentration (LC_50_), larvae of the pyrethroid-susceptible colonized strain *An. coluzzii* Ngousso were 16-fold more susceptible to chlorfenapyr than immature stages of another susceptible colony: *An. gambiae* Kisumu. Larval bioassays indicated 99.63 ± 0.2% mortality after 24 h at a discriminating concentration of 100 ng/ml in *Anopheles gambiae* and *An. coluzzii* larvae collected from seven locations in urban and rural areas of Yaoundé, Cameroon. By contrast, exposing emerging female adults from these populations to the recommended discriminating concentration (100 µg Active Ingredient (AI)/bottle) in bottle bioassays revealed variable mortality after 72 h, with values below the threshold of susceptibility (98%) in several tests. *Anopheles coluzzii* larvae and adults were fully susceptible, but mortality rates were slightly lower in *An. gambiae* adults compared to larvae (94 ± 1.5% vs 100%, Fisher’s exact test, *p* < 0.001). Piperonyl butoxide antagonized the activity of chlorphenapyr in *An. gambiae* adults. 100 ng/ml provides sufficient discriminative power for assessing the susceptibility of *An. gambiae* and *An. coluzzii* larvae to chlorfenapyr. Testing *An. gambiae* adults with 100 µg AI/bottle is likely to reveal inconsistent mortality values making it difficult to detect any emergence of resistance. Exploring different tests and accounting for variability between species are key to a reliable monitoring of *Anopheles* susceptibility to chlorfenapyr.

## Introduction

Long-lasting insecticidal nets (LLINs) and indoor residual spraying are the main vector-control tools used for malaria prevention (1,2). Chlorfenapyr (pyrolle) is an active ingredient in two new LLIN brands whose efficacy at reducing malaria incidence has been proven in field trials (3,4). Monitoring the susceptibility of wild populations to the active ingredient is vital for adopting proactive measures that will help preserve the efficacy of these new tools.

Standard bioassays are used to assess susceptibility by testing a discriminating concentration of chlorfenapyr on wild adult mosquito populations (5). Several studies have used dose-response assays to analyze the toxicity of chlorfenapyr and to derive discriminating doses that could be applied for monitoring the susceptibility of *Anopheles* female adults (6,7). In preparation for rollout of chlorfenapyr-containing LLINs, research programs coordinated by the US president’s malaria initiative (PMI) have used World Health Organization (WHO) bottle bioassays to test the susceptibility of laboratory colonies and field populations of *Anopheles gambiae* sensu lato in sixteen African countries (8). This investigation led to the conclusion that a concentration of 100-200 µg AI/bottle was suitable for susceptibility evaluation in wild anopheline populations. The WHO standard operating procedure for testing insecticide susceptibility of anopheline mosquitoes recommends using bottle bioassays while exposing *Anopheles* species to a discriminating concentration of 100 µg AI/bottle (5). Studies conducted so far have shown that although most populations are globally susceptible to 100 µg AI/bottle of chlorfenapyr, there are outliers that may require a higher dose (8–10).

Despite the availability of a standard procedure for testing *Anopheles* susceptibility to chlorfenapyr, it remains vital to explore supplementary techniques that may best reflect the toxicity mechanism of the active ingredient and improve our understanding of the potential selection pressures which could drive resistance (11). Chlorfenapyr’s mode of action differs remarkably from that of neurotoxic insecticides used for malaria vector control (12). This pyrrole acts as an uncloupler which inhibits oxidative phosphorylation in the mitochondrial membrane and impairs energy production within the cells leading to the death of the insect (13). Therefore, chlorfenapyr may induce lethal and sublethal effects that are only partially captured through 1-h exposure of adult populations to a dose of insecticide followed by the monitoring of short-term or delayed mortality as suggested by standard bioassays (5). Moreover, supplementary tests exploring alternative routes of exposure could help determine if outlier populations observed with standard bioassays are due to methodological artefacts or are subgroups exhibiting potential for resistance development.

Standardized procedures are also available for assessing the activity of synthetic chemicals on mosquito larvae, but the usefulness of this test for evaluating susceptibility to active ingredients applied against adults remains unexplored (14). However, it is well known that residual pesticide exposure at larval stages is a major driver of insecticide resistance observed in adult populations (15). Thus, testing susceptibility in *Anopheles* larvae could provide critical information for evaluating the efficacy of new malaria-control insecticides and the likelihood of resistance development. This approach has been informative for new insecticides such as neonicotinoids (16). It has been shown that monitoring mortality and some life table parameters in field-collected larvae reared in water containing a lethal concentration of a neonicotinoid could reveal patterns of survival and growth reflecting susceptibility profiles detected in adults using standard bioassays (17). Such investigations provided critical knowledge to support the growing evidence that residual pesticide exposure and cross-resistance contribute to the development of neonicotinoid resistance in *Anopheles* larvae and adults (17–22).

Available information on the use of agrochemicals in sub-Saharan Africa indicates that chlorfenapyr is not registered for use in crop protection in most African countries (23,24). Agricultural exposure is therefore unlikely to be a major source of selection pressure that could lead to resistance development. However, larval exposure to xenobiotics including herbicides, fungicides, insecticides, chemical fertilizers, and chemical pollutants is known to enhance insecticide resistance in adult mosquitoes (25–30). Notably, the development of chlorfenapyr resistance in insect pests is often associated with cross-resistance to other pesticides (31). Although multiple studies have detected no evidence of cross-resistance between chlorfenapyr and pyrethroids in *Anopheles* adults (6,9,32,33), the cross-reactivity of this active ingredient in larvae remains unknown. The rationale for assessing the toxicity of chlorfenapyr in *Anophele*s larvae is: (1) to test if xenobiotic exposure in aquatic habitats could enhance tolerance to the active ingredient and (2) to determine if larval and adult bioassays could be used concomitantly to better evaluate susceptibility in wild populations.

Here we used WHO guidelines for testing mosquito larvicides and WHO bottle bioassays to assess the susceptibility of two laboratory colonies and seven field-collected populations to chlorfenapyr. *Anopheles coluzzii* and *An. gambiae* larvae from Yaoundé were susceptible to 100 ng/ml of the active ingredient. Adults’ mortality was also high, but it was difficult to clearly ascertain the susceptibility profile of some wild *An. gambiae* populations using bottle bioassays and a discriminating concentration of 100 µg AI/bottle.

## Material and methods

### Ethics statement

Approval to conduct the study in the Centre region (N°: 1-140/L/MINSANTE/SG/RDPH-Ce), ethical clearance (N°: 1-141/CRERSH/2020) and research permit (N°: 000133/MINRESI/B00/C00/C10/C13) were granted by the Ministry of public health and the Ministry of scientific research and innovation of Cameroon.

### Study sites and mosquito populations

Mosquito larvae were collected from urban, sub-urban and rural areas of Yaoundé, the capital of Cameroon (Fig. 1). Seven locations were surveyed from February to December 2023. Natural reproduction sites of *Anopheles gambiae* sensu lato larvae were inspected, and immature stages were collected from standing waters using dippers and transported to the insectary. Larvae were identified as species using morphological keys (34,35). A subset of larvae were immediately tested in standard conditions (27°C, 80% relative humidity, light:dark = 12:12 h) upon arrival to the insectary. Another subset was fed with Tetramin® fish food and reared to adulthood. Emerging adult mosquitoes were provided with cottonwool pads dipped in 10% sugar solution until they were used for insecticide susceptibility tests. The two cryptic species, *An. gambiae* sensu stricto (hereafter referred to as *An. gambiae*) and *An. coluzzii*, are the dominant malaria vectors in Yaoundé (36). Both species were identified by subjecting 30 adult mosquitoes from each sampling site to a PCR technique enabling the discrimination of taxa of the *An. gambiae* complex based on mutations of the ribosomal DNA (37).

**Figure 1:**
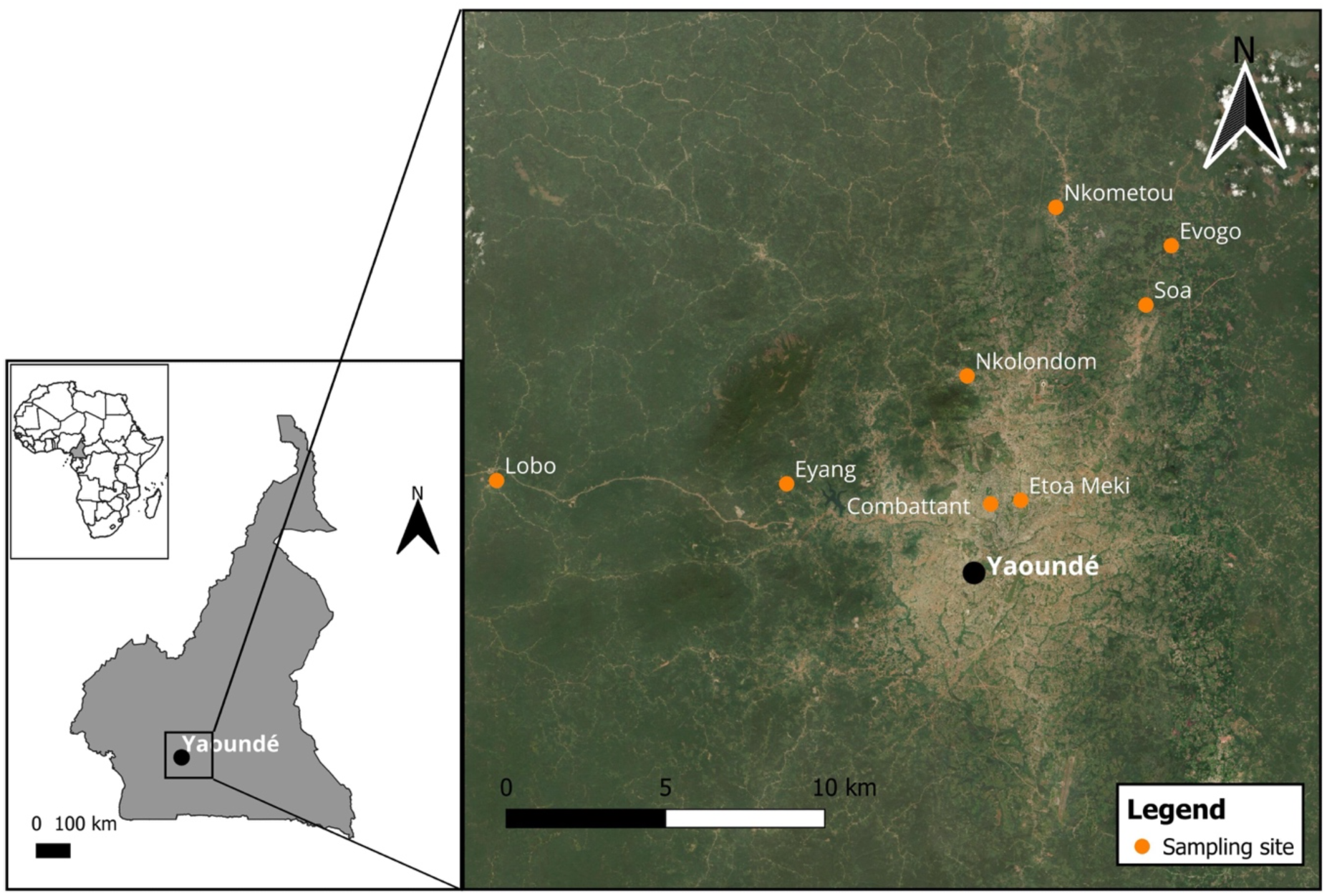
Map showing the sampling sites and the city of Yaoundé in the equatorial forest of Cameroon

### Insecticide

A technical-grade formulation of chlorfenapyr (PESTANAL®, analytical standard, Sigma-Aldrich, Dorset, United Kingdom) was tested. A stock solution used for larval bioassays and bottle tests was prepared by dissolving the active ingredient in absolute ethanol. The solution was prepared in 50 ml falcon tubes which were wrapped in aluminium foil and stored at 4 °C.

### Discriminating concentration determination and larvae susceptibility test

The lethal concentrations of chlorfenapyr were tested on two pyrethroid-susceptible colonized strains (*An. gambiae* Kisumu and *An. coluzzi* Ngousso) using concentration-response assays. Larval bioassays were conducted following WHO guidelines for testing mosquito larvicides (14). The protocol implemented in this study has been successfully used to test the susceptibility of *Anopheles* larvae to neonicotinoids and pyrethroids (17). Third instar larvae were exposed to increasing concentrations of the insecticide, and mortality was recorded at 24 h. Batches of 25 larvae were placed in 500-ml plastic trays filled with 200 ml borehole water containing the desired concentration of insecticide and covered with a net. Four replicates were tested for each insecticide concentration in addition to two controls without insecticide. The same volume of solvent (alcohol) used in tests was added to each control. After 24 h, larvae were considered dead if they were moribund and unable to move when touched with dropper. The following gradients were used to determine the lethal concentrations: *An. gambiae* Kisumu (1, 9, 20, 50, 70, 80, 100, 120 and 150 ng/ml); *An. coluzzi* Ngousso (1, 3, 5, 8, 20, 50 and 100 ng/ml). LC_50_ and LC_99_ corresponding to the concentrations needed to achieve 50 and 99% larval mortality, respectively, were measured at 24 h.

A discriminating concentration corresponding to LC_99_ was used to test the susceptibility of field-collected larvae to chlorfenapyr. The highest LC_99_ value between the two colonized strains *An. gambiae* Kisumu and *An. coluzzi* Ngousso was chosen. The larval population was considered susceptible if mortality was > 98% in water containing the discriminating concentration.

### Residual activity of chlorfenapyr in water

In this study, nominal concentrations were used without measuring the quantity of active ingredient dissolved in water. However, if chlorfenapyr is poorly soluble or instable in water, its concentration throughout the experiment could be different from the nominal value. We indirectly tested the relative stability of chlorfenapyr in water by monitoring the larvicidal activity of an LC_99_ concentration for seven consecutive days. Four batches of 25 filed-collected larvae were placed in 500-ml plastic trays filled with 200 ml water containing 100 ng/ml of chlorfenapyr. Controls consisted of two batches of 25 larvae placed in trays containing water and a volume of ethanol used in tests. Every 24 h, mortality was recorded, the solution was filtered to remove all larvae, alive or dead, and a new batch of 25 third instars was put in the tray. This process was repeated daily for seven days.

### Adult bioassays

The susceptibility of adult mosquitoes to chlorfenapyr was tested using the WHO bottle bioassays (5). 250-ml Wheaton bottles and their caps were coated with the insecticide following the Centers for Disease and Control (CDC) guidelines (38). The WHO recommended discriminating concentration (100 µg AI/bottle) was used. 24 h after coating bottles and caps with 1 ml of a solution containing the desired dose of chlorfenapyr, four replicates of 25 females (2–5 days old) were released into the test bottles and exposed for 1 h. After the exposure, mosquitoes were transferred into paper cups and provided with 10% sugar solution. Mortality was recorded at 72 h post-exposure. Two bottles coated with 1 ml of absolute ethanol were used as control. Based on a continent-wide study, wild *An. gambiae* sensu lato populations are susceptible to concentrations ranging from 100 to 200 µg AI/bottle (8). Due to this flexible range, mortality rates obtained with 100 µg AI/bottle in this study were not interpreted strictly based on WHO guidelines which imply that resistance is suggested when less than 98% of individuals die after the holding period (39). The susceptibility of female adults from a site located in the sub-urban area (Nkolondom) which displayed less than 98% mortality at 100 µg AI/bottle was tested for seven months.

The antagonistic effect of piperonyl butoxide (PBO) was tested using samples from the same site (Nkolondom). Female adults were pre-exposed to bottles coated with 4% PBO for 1 h before being released into test bottles treated with 100 µg AI/bottle of chlorfenapyr. After 1 h-exposure and 72 h-holding period, mortality was recorded and compared to the value obtained without PBO.

### Data analysis

Mortality values were presented at 24 h following the release of larvae in water containing the insecticide and at 72 h post-insecticide exposure for bottle bioassays using the mean and standard error. Abbot’s correction was not applied due to the control mortality never exceeding 10%. LC_50_ and LC_99_ values were calculated using a probit model implemented in the *ecotox* package in R version 4.2.2 (40). A ratio test was used to assess the difference in LC_50_ values between the two insecticide-susceptible colonies. Fisher’s exact test with a significance threshold set at 0.01 was applied to compare mortality rates between tests.

## Results

### Larval bioassays reveal susceptibility to chlorfenapyr

The concentration-response curves depicting the toxicity of chlorfenapyr for larvae of the pyrethroid-susceptible colonies *An. gambiae* Kisumu and *An. coluzzii* Ngousso are presented in Fig. 2, and the corresponding LC values are summarized in Table 1. LC_50_ was 16-fold higher in *An. gambiae* Kisumu larvae (22.9 ng/ml, 95% confidence interval (CI) [18.4-27.3]) compared to third instars of *An. coluzzii* Ngousso (6.3 ng/ml, [4.9-8.0]). A ratio test between LC_50_ values confirmed that the toxicity of chlorfenapyr was significantly higher in immature stages of *An. coluzzii* Ngousso than in *An. gambiae* Kisumu larvae (ratio test, p < 0.001). Also, LC_99_ obtained with *An. gambiae* Kisumu was 102 ng/ml [78.1-151] compared to 53 ng/ml [32-124] in *An. coluzzii* Ngousso. The highest LC_99_ (102 ng/ml) caused the maximum larvicidal effect (death of 100% of larvae) in both colonies. Therefore, 100 ng/ml was used as discriminating concentration to measure the susceptibility of wild mosquito larvae. Six field populations representing a total of 885 larvae were tested with 100 ng/ml. 100% mortality was observed within 24 h of dipping larvae in water containing the insecticide in all populations except one for which 24-h mortality was 97 ± 1% (Fig. 3). Two populations were identified as *An. coluzzii* (Combattant and Etoa Meki) by subjecting emerging adults to PCR. Emerging adults from Lobo, Eyang and Nklondom were 100% *An. gambiae*. Samples from the suburban areas, Soa, Nkometou and Evogo contained a mixture of *An. coluzzii* and *An. gambiae* at the following respective proportions: 30%:70%, 3%:97%, and 25%:75%. Larvae from Nkolondom were tested for four different months (May, September, October, and December 2023) and displayed 100% mortality at 100 ng/ml. The residual effect defined as the ability to maintain lethal activity towards a target organism for a certain amount of time was tested. Figure 4 shows the activity of chlorfenapyr over a period of seven days against larvae collected from Nkolondom. At 100 ng/ml, the active ingredient caused 100% mortality in batches of larvae tested every 24 h for seven days except the fourth day where mortality was 97 ± 1.7 %.

**Figure 2:**
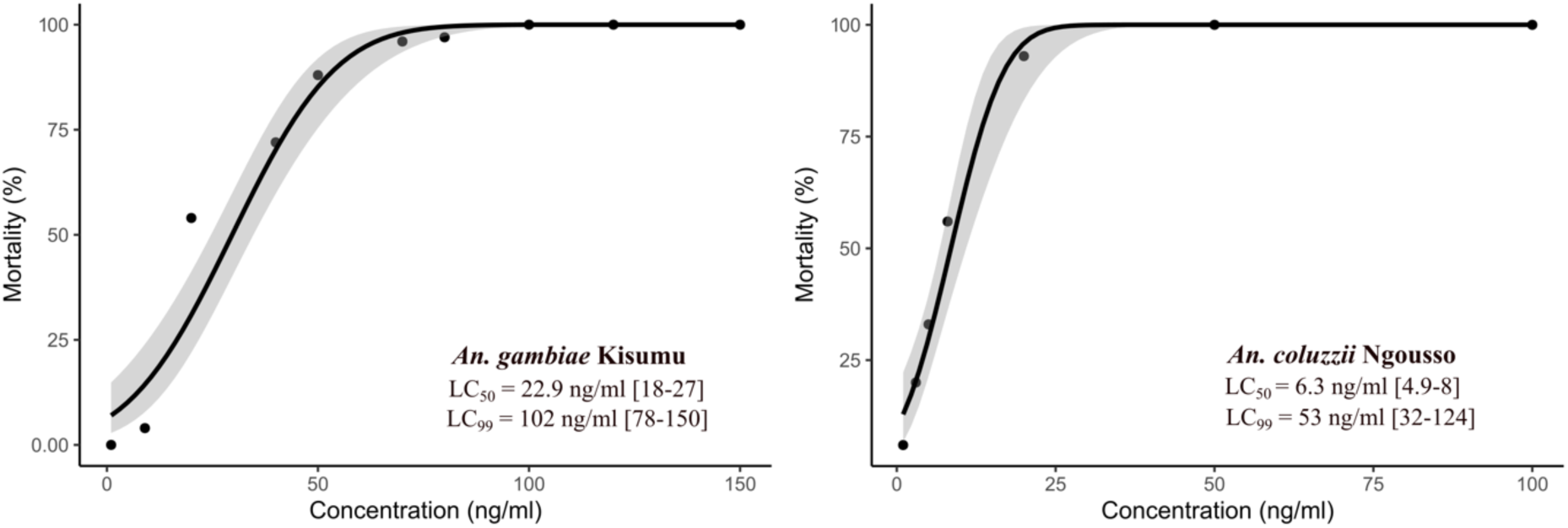
Concentration-response curves indicating the 24-h lethal concentrations of chlorfenapyr in third instar larvae. The grey band shows the standard error of the regression model. LC_50_ and LC_99_ for the pyrethroid-susceptible strains *An. coluzzii* Ngousso and *An. gambiae* Kisumu are shown with their 95% confidence intervals.

**Figure 3:**
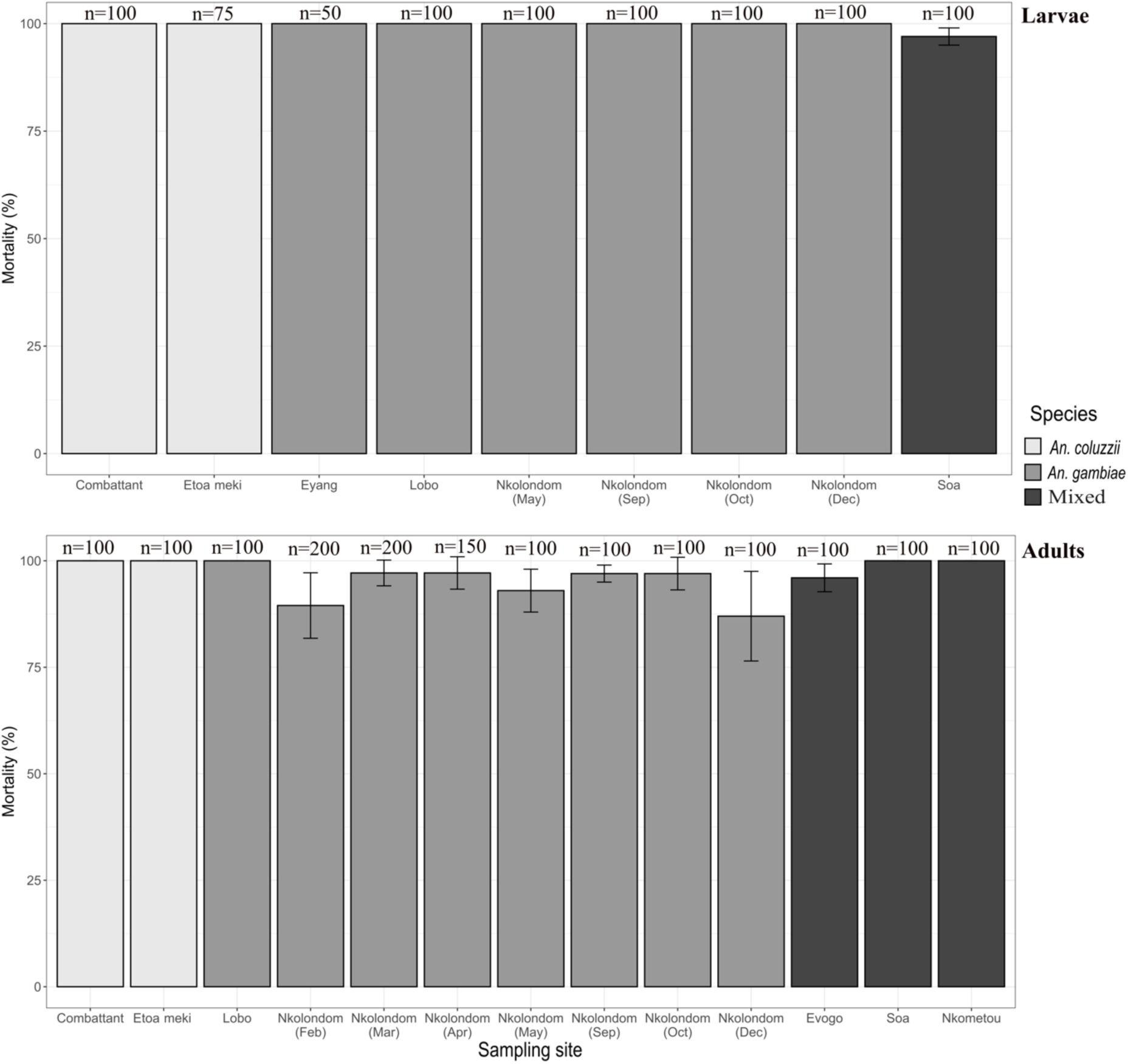
Mortality rates in field-collected *Anopheles coluzzii* and *An. gambiae* larvae and adults exposed to chlorfenapyr. Error bars represent the standard error of the mean.

**Figure 4:**
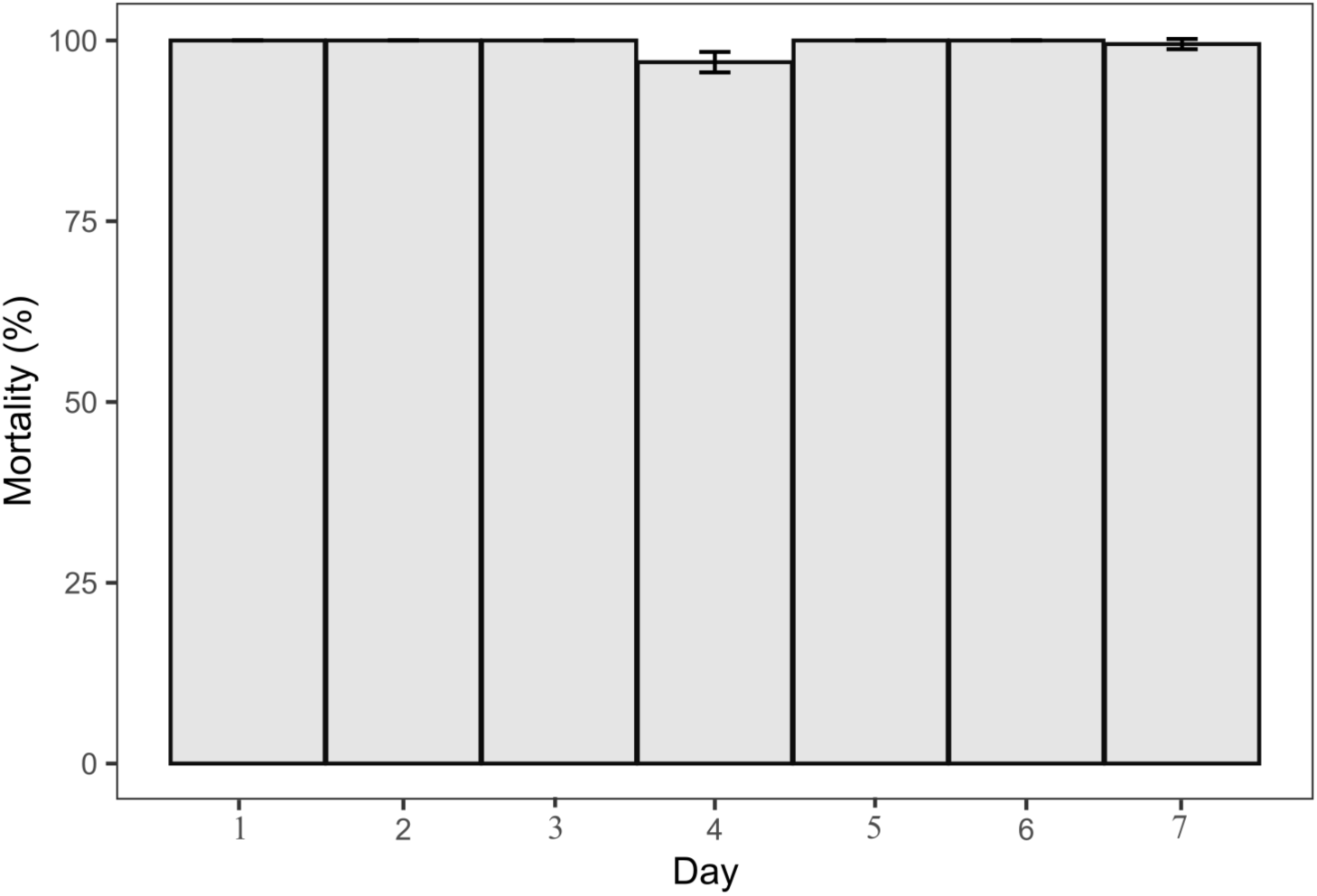
Residual activity of 100 ng/ml of chlorfenapyr monitored for seven consecutive days in field-collected *An. gambiae* larvae.

**Table 1:** Lethal concentrations, LC_50_ and LC_99_, of chlorfenapyr in third instar larvae estimated at 24 h post-exposure.

| Species | LC<sub>50</sub> value<br>(ng/ml) | 95% CI | Standard<br>error | LC<sub>99</sub> value<br>(ng/ml) | 95% CI | Standard<br>error | Chi-Square | p-value |
| --- | --- | --- | --- | --- | --- | --- | --- | --- |
| *An. gambiae*<br>Kisumu | 22.9 | 18-27 | 1.1 | 102 | 78-150 | 1.17 | 3.69 | < 0.001 |
| *An. coluzzii*<br>Ngousso | 6.3 | 4.9-8 | 1.13 | 53 | 32-124 | 1.38 | 3.02 | < 0.001 |

### Adult bioassays show variable mortality at 100 µg AI/bottle

At the discriminating concentration of 100 µg AI/bottle recommended for routine evaluation of *Anopheles* susceptibility to chlorfenapyr, both pyrethroid-susceptible colonies (*An. coluzzii* Ngousso and *An. gambiae* Kisumu) reached 100% within 72 h of holding period. However, variable mortality rates were observed in some field-collected populations (Fig. 3). Both *An. coluzzii* populations (Combattant and Etoa-meki) were susceptible, with 100% mortality recorded after 72 h. *An. gambiae* adults from Lobo and mixed populations from Soa and Nkometou were also fully susceptible. A slightly reduced mortality rate was observed in samples from Evogo (96 ± 1.6%). A longitudinal survey was conducted in Nkolondom, and the susceptibility profile of adult *An. gambiae* mosquitoes was monitored for seven months (February, March, April, May, September, October, and December 2023). Results showed a variation in mortality with the lowest values observed in December (87 ± 5.2%), February (91.5 ± 2.4%), and May (93 ± 2.5%) (Fig. 3). The antagonistic effect of PBO was assessed by pre-exposing 100 *An. gambiae* females from Nkolondom to 4% PBO before testing them with 100 µg AI/bottle. PBO significantly lowered mortality (from 91.5 ± 2.4% to 39 ± 17 %, Fisher’s exact test, *p* < 0.001, OR = 0.46, 95% CI = 0.31-0.67) indicating antagonism (Fig. 5).

**Figure 5:**
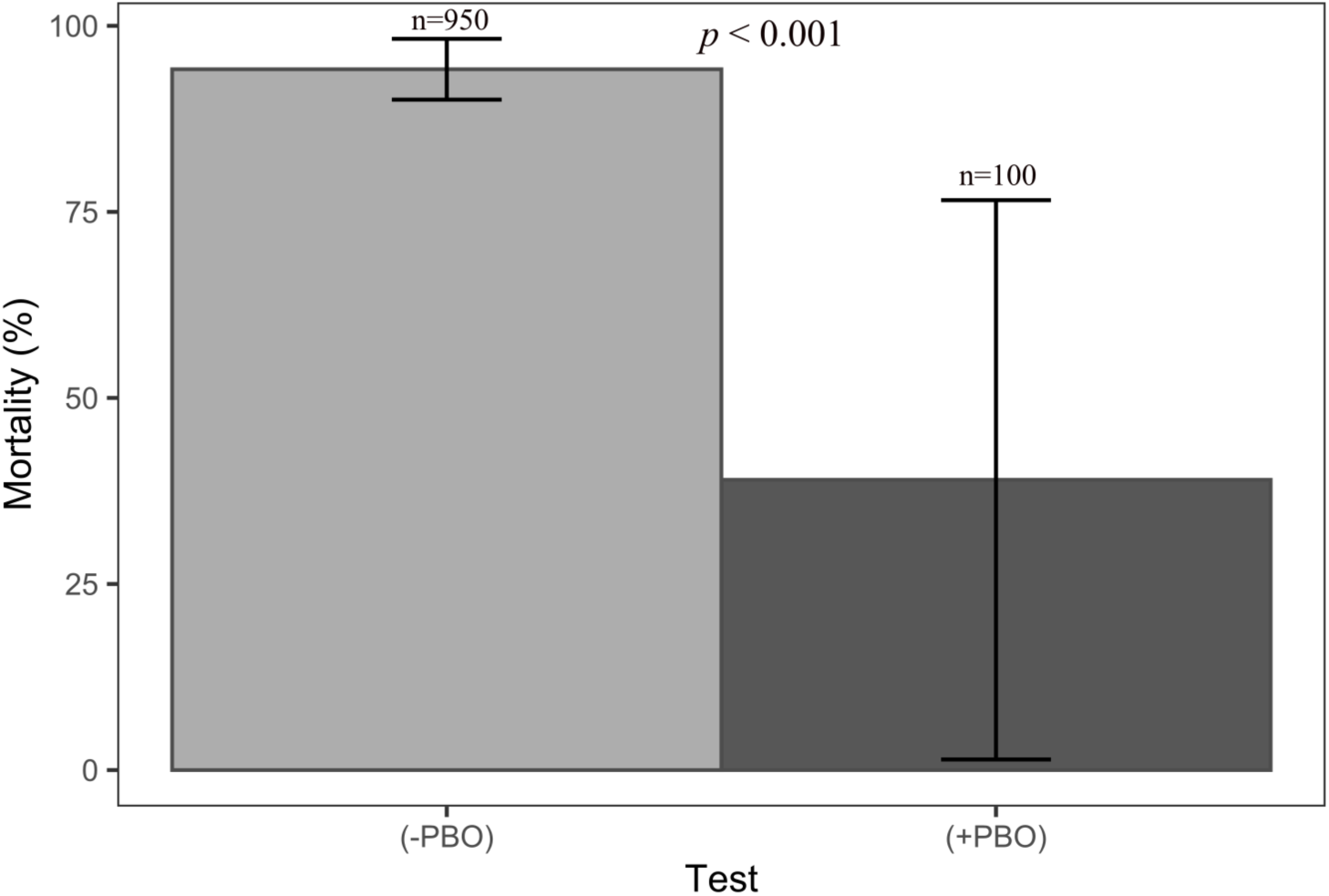
Antagonistic effect of PBO on the activity of chlorfenapyr. Mortality of *An. gambiae* female adults was significantly reduced within 72 h of insecticide exposure in the presence of PBO (Fisher’s exact test, *p* < 0.001).

### Mortality rates are consistent between larval and adult bioassays

Susceptibility profiles were compared between wild larvae (n = 825) and adult mosquitoes (n = 1550) (Fig. 6). It appeared that *An. coluzzii* larvae (n = 175) and adults (n = 200) displayed the same mortality rate: 100% after 24 h for larvae and 100% within 72 h of insecticide exposure for adults. In *An. gambiae*, mortality was slightly lower in adults (94 ± 1.5%) (n = 1050) compared to larvae (100%) (n = 550) (Fisher’s exact test, *p* < 0.001, OR = 0.09, 95% CI = 0.019-0.3) (Fig. 6). Mortality was not significantly different between the mixed larval population from Soa (97 ± 1%) (n = 100) and mixed adults from Evogo, Soa and Nkometou (98.6 ± 0.7%) (n = 300) (Fisher’s exact test, *p* = 0.37, OR = 2.24, 95% CI = 0.32-13.51).

**Figure 6:**
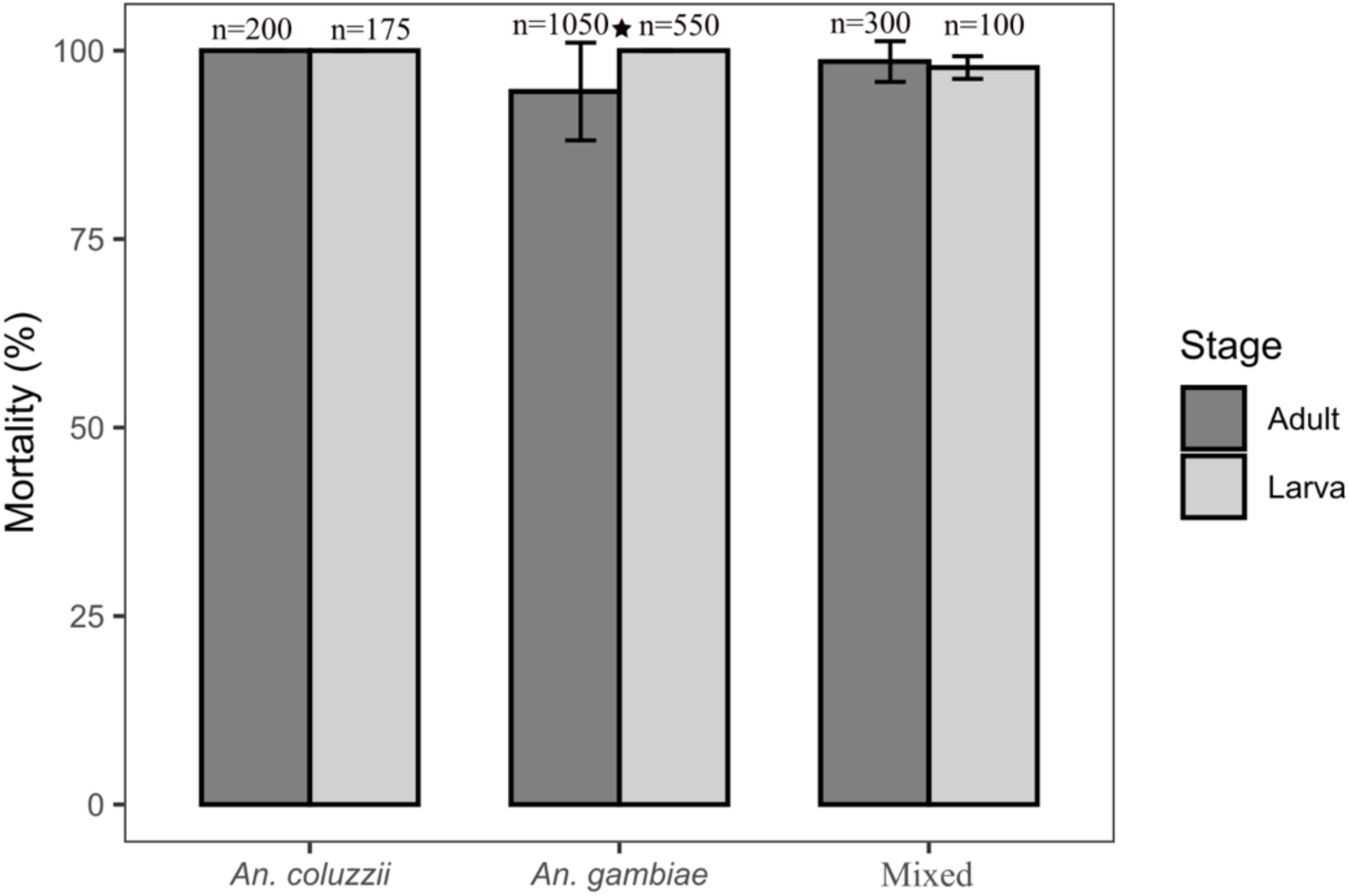
Comparison of mortality rates between larvae exposed to 100 ng/ml of chlorfenapyr for 24 h and adults exposed to 100 µg AI/bottle for 1 h and monitored for 72 h. * Fisher’s exact test indicates slightly lower mortality in *An. gambiae* adults compared to larvae (*p* < 0.001)

## Discussion

The toxicity of chlorfenapyr has been tested on immature stages of various insect pests using diverse routes of exposure (41–44). Regarding mosquitoes, Yuan et al, 2015 (45) measured a median lethal concentration (LC_50_) of 35 ng/ml after 24 h by dipping early fourth instar larvae of *Culex pipiens pallens* in 200 ml water containing a concentration of chlorfenapyr. In the present study, the larvicidal toxicity of chlorfenapyr was evaluated in *Anopheles* mosquitoes using an experimental setup comparable to Yuan et al’s protocol. Based on persistent exposure of third instar larvae in water containing the active ingredient, it was observed that LC_50_ for the pyrethroid-susceptible strain *An. gambiae* Kisumu (22.9 ng/ml) was within the range of values detected when chlorfenapyr toxicity was measured in *Culex pipiens pallens*. LC_50_ for the *An. coluzzii* strain Ngousso was significantly lower (6.3 ng/ml). Studies have revealed that the toxicity of other new vector-control insecticides such as neonicotinoids was also higher in *An. coluzzii* larvae compared to immature stages of *An. gambiae* (17). Chlorfenapyr was 16 times more toxic to *An. coluzzii* Ngousso than it was to *An. gambiae* Kisumu which highlights the need for considering variability between species when implementing susceptibility tests for this active ingredient (5,8,46).

Approximately 100% larval mortality was observed after 24 h in all laboratory colonies and field populations tested using a discriminating concentration of 100 ng/ml which indicates that chlorfenapyr has high efficacy against *Anopheles* larvae. Comparatively, *An. gambiae* third instars collected from one of the study sites (Nkolondom) display less than 10% mortality after 24 h when they are reared in water containing a lethal concentration of deltamethrin, clothianidin, imidacloprid or acetamiprid (17). The higher efficacy of chlorfenapyr on immature stages of *Anopheles* mosquitoes compared to clothianidin has also been demonstrated by resistance selection in laboratory conditions (18). The experiment revealed that after ten generations of resistance selection, both selected larvae and selected adults of a pyrethroid-resistant *An. gambiae* colony remained susceptible to chlorfenapyr while developing resistance to clothianidin (18). Although the findings have yet to be validated in other geographic contexts, the susceptibility of *An. gambiae* and *An. coluzzii* larvae from Yaoundé indicates a lack of cross-resistance between chlorfenapyr, pyrethroids and neonicotinoids. For instance, larvae from Nkolondom which are susceptible to chlorfenapyr are highly resistant to deltamethrin, acetamiprid, imidacloprid and clothianidin (17). Studies have also highlighted a lack of cross-reactivity between chlorfenapyr, pyrethroids, organophosphates and carbamates in *Anopheles* adults (6,32,33).

Adult bioassays revealed variable mortality when the recommended discriminating concentration of 100 µg AI/bottle was used. Oxborough et al, 2021 (8) also observed dose-dependent susceptibility variations in some *An. gambiae* sensu lato populations. Although susceptible colonies were killed with concentrations between 10 and 50 µg AI/bottle in bottle bioassays, 100 to 200 µg AI/bottle were needed to reach 100% mortality in most wild adult populations. Median mortality was > 99% when 100 µg AI/bottle was used, but some outlier populations exhibited mortality rates below 98% at this concentration (8). A recent investigation also reported mortality rates below 98% in several *An. gambiae* s.l. populations exposed to 100 µg AI/bottle (9). In the present study, we have detected variations with mortality < 98% within 72 h of insecticide exposure in monthly tests conducted on *An. gambiae* populations from Nkolondom. Although seasonal changes can account for such variability, another possible explanation is that 100 µg AI/bottle of chlorfenapyr may not be sufficient for monitoring susceptibility in wild *An. gambiae* populations as initially suggested by Oxborough et al (8). Further studies are needed to confirm if higher doses should be recommended for testing some *Anopheles* species.

When adult mosquitoes from Nkolondom were pre-exposed to PBO before being tested with 100 µg AI/bottle of chlorfenapyr, a significant reduction in mortality was recorded. This antagonistic effect of PBO on the efficacy of chlorfenapyr has been documented in insect pests (31,45). Chlorfenapyr is a proinsecticide whose transformation into a toxic metabolite (tralopyril) is catalysed by cytochrome P 450 enzymes (CYPs), hence the reduction in activity when CYPs are inhibited by PBO (12,13,47,48). This finding suggests that downregulation of some CYPs may be a potential mechanism *Anopheles* could exploit to develop resistance to chlorfenapyr as is the case in the two-spotted spider mite, *Tetranychus urticae* (49).

Because of the difference in the modes of exposure, it is difficult to directly compare susceptibility profiles revealed by larvicidal activity testing and adult bioassays. Indeed, larval tests were based on persistent exposure for 24 h while mosquitoes were challenged to the insecticide for only 1 h in adult bioassays. However, in *Culex pipiens pallens* for example, it has been shown that chlorfenapyr toxicity followed the same trend in larvae dipped in water containing the insecticide for 24 h and in adults tested using the topical application method and WHO impregnated papers (45). Moreover, evidence from chlorfenapyr resistance selection in *An. gambiae* indicated the same susceptibility level in selected larvae and in selected adults (18). Despite the potential bias that may have been introduced by the difference in modes of exposure, our results revealed that larval and adult susceptibility mirrored each other and that larval and adult bioassays could be used concomitantly to assess the efficacy of chlorfenapyr in *Anopheles* mosquitoes. *An. gambiae* and *An. coluzzii* populations sampled from Yaoundé were susceptible to chlorfenapyr as revealed both by larval and adult bioassays. As mentioned previously, some tests conducted on adults indicated mortality values below the threshold of susceptibility (98%), but this result was not interpreted as evidence of resistance. Testing higher concentrations of the active ingredient and exploring alternative routes of exposure including topical application or feeding assays used in other insect pests could shed light on the status of outlier populations that are being detected with standard adult bioassays (31,45).

The larvicidal activity of chlorfenapyr in water was confirmed for seven consecutive days, which implies that the active ingredient was soluble and stable in water. However, direct chemical measurements are needed to detect the concentrations of chlorfenapyr in aqueous solutions. Moreover, additional tests on susceptible colonies and field populations of different malaria vector species have yet to be conducted to derive discriminations concentrations that could be applied for susceptibility monitoring using larval bioassays. However, the lethal concentrations detected in this study provided satisfactory discriminating power for screening populations of the sibling species *An. gambiae* and *An. coluzzii*.

## Conclusions

Larval bioassays can be used to test the susceptibility of anopheline mosquitoes to chlorfenapyr. Conducting both larval and adult bioassays could help disentangle the role of candidate selection pressures such as agricultural pesticide use and cross-resistance that may affect susceptibility. If tolerance to chlorfenapyr emerges in *Anopheles* species, screening both larvae and adult mosquitoes could facilitate a more comprehensive appraisal of the genetic basis of resistance.

## Acknowledgments

We thank farmers from Nkolondom for granting us access to the farmland.

## Funding

This study was supported by a National Institutes of Health grant (R01AI150529) to CK.

## Notes

### Competing Interest Statement

The authors have declared no competing interest.

